# Trace Elements in Fish: Assessment of bioaccumulation and associated health risks

**DOI:** 10.1101/2024.09.08.611911

**Authors:** Saima Naz, Qudrat Ullah, Dalia Fouad, Abdul Qadeer, Maria Lateef, Muhammad Waqar Hassan, Ahmad Manan Mustafa Chatha

## Abstract

Industrial and agricultural water run-off are polluting the aquatic ecosystem by depositing different toxic trace elements (TTEs) in riverine system. It has become a global concern impacting not only the well-being of aquatic organisms but human health as well. Current study evaluated the impact of four TTEs (Cadmium (Cd), copper (Cu), lead (Pb), and nickel (Ni)) in three organs (liver, gills, and muscles) of five fish species viz, *Rita rita, Sperata sarwari, Wallago attu, Mastacembelus armatus,* and *Cirrhinus mrigala* collected from right and left banks of Punjnad headworks during winter, spring and summer. We investigated accumulation (mg/kg) of these TTEs in fish in addition to the human health risk assessment by estimating exposure hazards, hazardous index (THQ and TTHQ) and metal pollution index (MPI). The obtained results showed that *W. attu* accumulated significantly more TTEs (p < 0.00) as compared to other fish. Among seasons, summer had significantly more (p < 0.00) accumulation of TTEs than other seasons. Lead (Pb) accumulation was highest across TTEs in fish liver as compared to gills and muscles. Right bank showed higher accumulation (p < 0.00) of all TTEs in all fish species in contrast to the left bank. The human health risk assessment showed that Cd and Pb had higher exposure levels than Cu and Ni. Furthermore, the THQ was in the order of Cd > Pb > Ni > Cu. All fish species had THQ 1 for Cd and Pb and TTHQ > 1 for all fish. MPI index showed moderate to high level of TTE contamination if all fish species. The study concluded that right bank has higher metal accumulation than left bank. However, fish consumption from both of the study site was not safe for human consumption.

## 1. Introduction

The physical, chemical, and biological characteristics of water, typically in relation to its suitability for a specific purpose, such as drinking, swimming, agriculture, or supporting aquatic life is referred as Water quality (1, 2). It is measured by the levels of organic and inorganic chemicals in the water, along with certain physical properties (3). Natural water can become contaminated by untreated waste from industries, agriculture, and technology, which often contain metallic compounds in traces (4). These trace elements are particularly harmful because they do not break down naturally, (5, 6), can build up in food chains (7), and can disrupt aquatic ecosystems and the organisms living in them thus become toxic for living organisms (8). These toxic trace elements (TTEs) enter into water through waste from industries like tanneries, textiles, metal finishing, mining, dyeing, ceramics, and pharmaceuticals. When fish absorb TTEs, they can pass them on to humans through the food chain, leading to serious health risks, including potentially fatal consequences (6, 9, 10).

The accumulation of TTEs in fish mainly depends upon the concentration of these metals in the aquatic environment and the duration of exposure. However, several other factors, such as pH, water salinity, hardness, temperature, fish size and age, ecological needs, life cycle, capture season, and feeding habits, also significantly influence metal accumulation (8). When metal levels in fish tissues become excessively high, they can become toxic (11, 12). Toxic trace elements are stable and persistent contaminants in aquatic environments. Some trace elements, like zinc (Zn), copper (Cu), iron (Fe), and manganese (Mn), are essential for metabolic processes in organisms but exist in a narrow range between being beneficial and toxic. Other trace elements such as cadmium (Cd), mercury (Hg), chromium (Cr), and lead (Pb), can be extremely toxic even at low concentrations, especially under certain conditions, making regular monitoring of sensitive aquatic environments necessary. The process of bio-magnification can cause pollutants to reach toxic levels in species higher up the food chain, particularly in freshwater systems (13, 14).

Various fish species are commonly used to assess the health of aquatic life and their ecosystems, especially as the pollution caused by TTEs continues to rise (15). Different biological techniques are employed to evaluate the toxic levels of metals and their impact on the behavior, physiological sensitivity, and morphological characteristics of fish and other organisms (16). Fish, which occupy a critical position at the top of the aquatic food chain, are particularly sensitive to pollution caused by TTEs. For instance, Prolonged exposure to Cd can lead to its accumulation in various tissues of fish, affecting the structure and function of vital organs like the gills, liver, and gonads (17). The use of cadmium-containing fertilizers, agricultural chemicals, pesticides, and sewage sludge on farmland can contribute to water contamination (18). Likewise, When fish are exposed to Copper (Cu), notable changes in the liver include hepatocyte vacuolization, necrosis, shrinkage, nuclear pyknosis, and an increase in sinusoidal spaces were noticed (19). Copper is highly toxic in aquatic environments, affecting fish, invertebrates, and amphibians, with all three groups liver, even at low environmental concentrations (20). In fish it tends to accumulates in various organs of fish like liver, kidney, gills, bones, fins, and muscles (21, 22). In Cu-exposed fish, distinct changes were observed in the liver include hepatocyte vacuolization, necrosis, shrinkage, nuclear pyknosis, and an increase in sinusoidal spaces (19). Trace elements are primarily absorbed by fish through their gills, but uptake can also occur through the intestinal epithelium (23). In fish species such as *Oreochromis niloticus* and *Lates niloticus*, exposure to TTEs has led to severe degenerative and necrotic changes in their intestinal mucosa (24). Lead (Pb) is a toxic trace element that causes a wide range of adverse health effects, which vary depending on the dose. Both fish and humans are primarily exposed to lead through ingestion and inhalation. Lead tends to accumulate in muscles, bones, blood, and fat, making it a potent environmental pollutant with significant implications for human health (25, 26). Nickel (Ni) is another trace metal widely distributed in the environment, released from both natural sources and human activities, including stationary and mobile sources. It can be found in the air, water, soil, and biological materials (27, 28). In fish, exposure to high concentrations of Ni has been linked to higher mortality rates. For showing similar sensitivity to chronic toxicity.

The Mrigal fish like *Cirrhinus mrigala* are widely used as food and holds significant economic importance in its native regions (29). *Wallago attu*, a large freshwater catfish, is found across rivers, reservoirs, and connected watersheds in the Indian subcontinent, including India, Pakistan, Bangladesh, Nepal, Burma, Sri Lanka, and parts of Southeast Asia like Thailand, Vietnam, Cambodia, and Indonesia (30). Its rapid growth, elongated silvery body, and high nutritional quality have made it a focus of aquaculture development. However, the declining wild populations of *Wallago attu* have led to its classification as an endangered species (31) . Increased consumer demand has driven the development of intensive aquaculture for this species in Asian countries (32). The spiny eel, *Mastacembelus armatus*, is one of the most common and economically significant inland teleost species in Asia, known for its high market and nutritional value (33). It is a popular table fish due to its delicious taste and high nutritional content, and it is also popular as an aquarium fish. Recently, it has gained attention as an indigenous ornamental fish exported from India to other countries (34). The Indus catfish, *Sperata sarwari,* naturally inhabits a variety of freshwater bodies across South Asia, from Afghanistan to Thailand (35). It primarily lives in riverine habitats but can also survive and breed in ponds, lakes, tanks, channels, and reservoirs (34). *Sperata sarwari* is valued for human consumption and is known for its fighting ability when hooked. *Rita rita*, a freshwater catfish of the family Bagridae, inhabits tropical rivers and estuaries and is an important food fish with high nutritional value. It is also used as a species of choice for monitoring riverine pollution (36). Known for its hardiness and tolerance to wide fluctuations in water quality due to human activities, *Rita rita* has recently become a key species in aquatic pollution monitoring and biomarker response studies (36).

Punjand The present topic has long been neglected in Pakistan as only a few studies have reported on genetic diversity and identification of fishes in River Punjand (37). However, to date no scientific literature is available on the effects of TTEs on fish or other aquatic organisms in RB and LB of Punjand headworks. In order to fill this gap, present study is specifically designed to study the effects of four TTEs (Cd, Cu, Ob, and Ni) being more toxic and in higher concentrations (37) on LB and RB of Punjnad headworks. Current study explored the accumulation of selected TTEs in three organs (liver, gills, and muscles) of five fish species (*R. rita*, *S. sarwari*, *W. attu*, *M. armatus*, and *C. mrigala*, during winter, spring and summer collected from Left and Right banks of Punjnad headworks. Furthermore, the health risk assessment of these TTEs to human health was also determined to explore its toxic effects on mature and juvenile consumers. The results of this study would be helpful in future for the health risk assessment in different species of all the two stated sites of River Punjand and remedies for lowering the pollution.

## 2. Materials and Methods

### 2.1 Study Area

The Punjnad headworks, located near Uch Sharif in Punjab, Pakistan, is a crucial agricultural region where the five rivers (Beas, Sutlej, Ravi, Chenab, and Jhelum) of Punjab merge into one river namely Chenab. This area is vital for meeting the irrigation demands of the various districts of Bahawalpur and Rahim Yar Khan, as well as the northern parts of Sindh. It is characterized by extensive industrial and agricultural activities, which significantly contribute in the accumulation of various TTEs in river Chenab and Punjnad headworks. Current study was conducted at left bank (LB) (Latitude (28°57’06.3”N); Longitude (70°30’32.8”E)), and right bank (RB) (Latitude (28°56’45.4”N); Longitude (70°29’49.7”E)), Uch Sharif, Punjab, Pakistan (**Figure 1**). The river banks were selected as no previous work has been reported on the accumulation of TTEs at these sites. Sampling of five fish species viz*, Rita rita, Sperata sarwari, Wallago attu, Mastacembelus armatus,* and *Cirrhinus mrigala* was carried once a month from November, 2021 to July, 2022. These sampling months were divided into three sampling seasons (winter, spring, and summer). The research specifically focuses on riverbanks because considering seasonal variations in trace metals is crucial for a comprehensive investigation. Contaminated irrigation water can significantly impact crops grown along the riverbanks, posing serious risks to agricultural productivity and food safety. The experiment was designed to check the deleterious effects of nickel, chromium, copper and lead in organs (liver, gills and Muscles) on five freshwater fish species.

**Figure 1:**
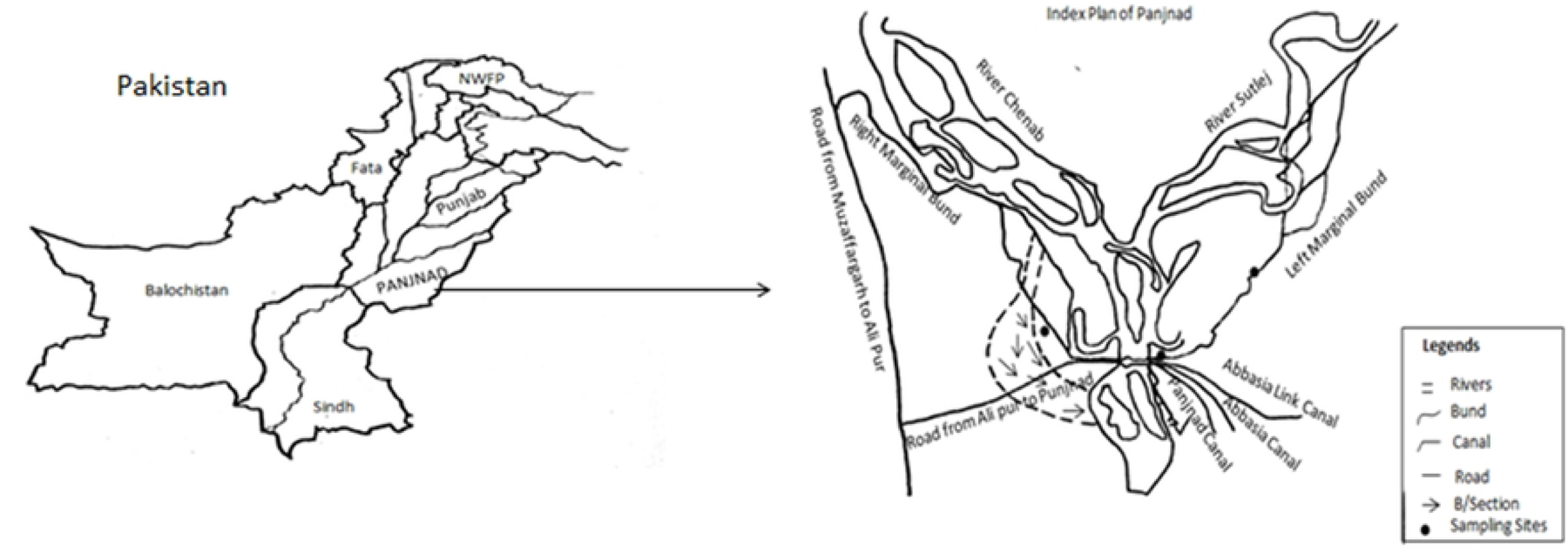
Study area at right bank (RB) and left bank (LB) of Punjand headworks, Uch Sharif, Punjab, Pakistan. The Sampling sites are indicated by (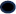).

### 2.2 Fish Sampling

Samples from each species were randomly collected one a month during day from study sites using a gauze net measuring 100 m × 6 m, with a mesh size of 60 mm. Sampling was conducted across three different seasons: winter, spring, and summer. After collection of samples were placed inside labeled polythene bags, the fish were immediately stored in a storage box (Coleman 48 Quart icebox) with crushed dry ice to keep the samples fresh and then transported to the Zoology Laboratory at Government Sadiq College Women University, Bahawalpur. The average weight (g), average fork length (inches), and average total length (inches) of the fish were recorded for all samples fish specimen (**Table S1**). In the laboratory, the samples were stored at −20°C in a freezer (Haier −25°C biomedical) until further analysis. The initial identification and common names of the fish species were verified with the assistance of local fish catchers and sellers. An expert taxonomist used systematic keys to accurately identify all species and correct any misidentifications (38).

### 2.3 Acid Digestion

The acid digesting technique was utilized to evaluate the heavy metal concentration in the gills, liver, and muscles of selected fish (39). For the analysis, fish organs were isolated and digested using a mixture of analytical grade HNO_3_ (65% - SIGMA-ALDRICH) and HClO_4_ (60% - DAEJUNG) in a 3:1 volume ratio. The digestion process followed the Clesceri, Eaton (40) method, lasting 4–5 hours, and was carried out on a hot plate (ANEX Deluxe Hot Plate AG-2166-EX) at 100°C. After digestion, the samples were allowed to cool to room temperature (25°C) and then filtered through Whatman No. 42 filter paper. The filtered samples were stored in sealed bottles and sent to the Central Laboratory of Muhammad Nawaz Shareef University of Agriculture (MNSUA) in Multan for detection TTEs (Cd, Cu, Pb, Ni). Following digestion, the concentrations of TTEs were determined using an acetylene air flame Spectroscopy using Atomic Absorption Spectrophotometer (AAS) (Analytik Jena: NovAA 400 P). The samples were subsequently tested regarding the content of TTEs in accordance with the prescribed instrument settings including specific limit of detection (**Table 1**). A blank was generated for every sample, and modifications were made using reference to the blank to ensure reproducibility and quality control of the analysis performed at Central Laboratory, MNSUA. The accuracy and precision of the tests were verified by comparing the results with the reference material (CRM IAEA 407) supplied by the International Atomic Energy Agency (IAEA). Additionally, the analysis of blanks and standards demonstrated satisfactory performance in heavy metal determination, with recoveries ranging from 95% to 101% for the metals examined.

**Table 1.** Condition of atomic absorption spectrometer used for the detection of heavy metal concentration.

| Metals | Wavelength (nm) | Gas | Support |
| --- | --- | --- | --- |
| Cadmium (Cd) | 228.8 | Acetylene | Air |
| Iron (Fe) | 248.33 | Acetylene | Air |
| Lead (Pb) | 217 | Acetylene | Air |
| Nickel (Ni) | 232 | Acetylene | Air |
| Zinc (Zn) | 213.86 | Acetylene | Air |

### 2.4 Health risk assessment

Human health risk assessment of toxic trace elements (TTEs) in fish muscles is important to highlight the bioaccumulation and harmful effects of trace elements like Cd, Cu, Pb, and Ni, which can pose serious health risks to consumers. Understanding the levels of TTEs in fish helps in identifying potential hazards, guiding regulatory limits, and ensuring food safety, thereby protecting public health from long-term exposure to toxic substances. Current study considered two main categories (regular and seasonal) of fish consumers. Regular consumers, eat fish on daily basis for the whole year. These are mostly fisherman or the population which lives very close to the rivers and have easy and cheap access to freshwater fish. Seasonal consumers, eat fish only during cooler months and avoid eating fish during hot weather. These two categories were further divided into two more categories (mature and juvenile). Mature consumers eat higher quantity of fish and have a higher body weight (kg), while juvenile consumers eat lower portion of fish and have comparatively lower body weight. The human health risk assessment of carried out based a few mathematical equations. These equations used a few constants to estimate the risks associated with the consumption of fish having toxicity from selected TTEs (**Table 2**).

#### 2.4.1 Ingestion exposure estimation

The estimation of ingestion exposure 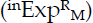 of selected TTEs in selected categories was calculated based on the following equations (**Equation 1-4**) (2).

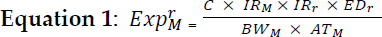

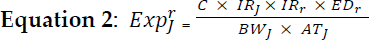

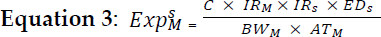

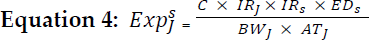

where 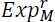 is the ingestion exposure of TTEs in mature regular consumers, 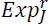 is the ingestion exposure of TTEs in juvenile regular consumers, 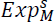 is the ingestion exposure of TTEs in mature seasonal consumers, and 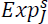 is the ingestion exposure of TTEs in juvenile seasonal consumers.

### 2.4.2 Target hazardous quotients estimation

Target hazardous quotients (THQs) are utilized to assess the potential non-carcinogenic health risks associated with exposure to TTEs present in the edible parts of fish (muscles), in accordance with the health risk assessment guidelines provided by the USEPA (41). The THQs dues to fish consumption, for the regular and seasonal (mature and juvenile) consumers was calculated using following equations (**Equation 5-8**).

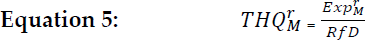

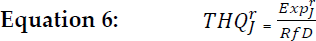

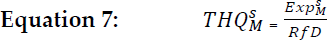

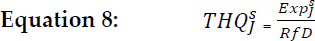

THQ represents the Hazard Quotient, calculated based on ingestion at the corresponding exposure level; RfD represents the reference dose for the potential hazardous health effects caused by contaminants through ingestion of TTEs. Current study used RfD value of 0.001, 0.04, 0.004, and 0.02 for Cd, Cu, Pb, and Ni respectively. THQ > 1 represent the potential toxic effects of TTEs to human consumption while THQ < 1 represent that fish is safe for human consumption.

#### 2.4.3 Total Target hazardous quotient estimation

The Total target hazardous quotient (TTHQ) is the sum of THQs for individual TTE and represent the cumulative exposure effects of all TTEs to human health. Like THQ, the TTHQ > 1 represent the potential toxic effects of TTEs to human consumption while TTHQ < 1 represent that fish is safe for human consumption (2). The TTHQs dues to fish consumption, for the regular and seasonal (mature and juvenile) consumers was calculated using following equations (**Equation 9-12**).

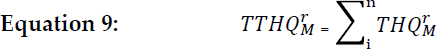

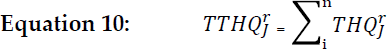

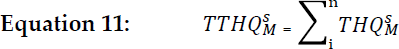

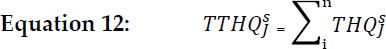

#### 2.4.4 Meral pollution index (MPI)

Metal pollution index of TTEs (MPI_TTE_) is determined using the following equation (42, 43):

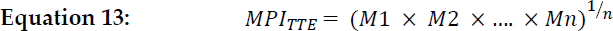

where *M*1 is the concentration (mg/kg) of the first TTE, *M*2 is the concentration of the second TTE and Mn is the concentration of the *nth* TTE in the muscle of fish, while *n* is the number of TTE studied.

### 2.5 Statistical analysis: -

The data collected in this research are presented as mean ± S.E. Normal distribution was observed within each group, and statistical analysis was conducted using one-way analysis of variance (ANOVA) with IBM SPSS Statistics software (version 20). Post hoc Duncan multiple range test was employed to determine differences in mean values with *P* < 0.05. Furthermore, the data were also subjected to

Pearson’s correlation analysis using Minitab (Version: 19.1.1)) to determine the correlation between various fish species for the accumulation of TTEs. We also applied Principal component analysis (PCA), and hierarchical cluster analysis (HCA) by using PAST (version: 4.03) to explore the relationship between fish species and studied TTEs.

**Table 2.** Different terms with definitions and values are used for the health risk assessment of toxic trace elements in fish muscles.

| Term used | Definition | Value |
| --- | --- | --- |
| $IR_M$ | Ingestion rate of mature person | 0.20kg/day |
| $IR_J$ | Ingestion rate of juvenile person | 0.10kg/day |
| $EF_s$ | Exposure frequency | 150 days/year |
| $EF_r$ | Exposure frequency | 365 days/year |
| $ED_M$ | Exposure duration of mature person | 30 years |
| $ED_J$ | Exposure of juvenile person | 12 years |
| $BW_M$ | Body weight of mature person | 70 kg |
| $BW_J$ | Body weight of juvenile person | 30 kg |
| $AT_M$ | Average days (Exposure) of mature person | 10,950 days |
| $AT_J$ | Average days (Exposure) of juvenila person | 4380 days |

## 3. Results

### 3.1 Accumulation of toxic trace elements (TTEs) in different organs of fish

A study was conducted to assess the accumulation of four TTEs (Cd, Cu, Pb, and Ni) in three different organs (liver, gills, and muscles) in five fish species (*R. rita*, *S. sarwari*, *W. attu*, *M. armatus*, and *C. mrigala*) during three different seasons (winter, spring, and summer) from right (RB) and the left banks. The season wise analysis of accumulation (mg/kg) of TTEs showed that during winter, across all species and TTEs, the liver consistently showed the highest concentration of TTEs, followed by gills and muscles. This trend is expected because the liver is a primary detoxifying organ, where metals accumulate more prominently. *Wallago attu* consistently showed the highest accumulation among all metals and organs compared to other species, particularly in the liver, where it has the highest recorded values for Cd (10.12±2.18 in LB and 15.25±2.47 in RB), Cu (5.65±1.57 in LB and 7.48±1.81 in RB), Pb (41.61±4.99 in LB and 46.03±10.56 in RB), and Ni (8.94±2.21 in LB and 11.68±1.23 in RB). On the other hand, *C. mrigala* has the lowest accumulation TTEs having the significantly lower accumulation (Cd (1.38±0.15 in LB and 2.08±0.52 in RB), Cu (0.57±0.06 in LB and 1.64±0.21 in RB), Pb (8.29±1.65 in LB and 11.87±1.90 in RB), and Ni (1.48±0.24 in LB and 2.39±0.26 in RB) in fish muscles. Furthermore, the pattern of TTEs’ accumulation in fish species was in the order of *W. attu* > *M. armatus* > *S. sarwari* > *R. rita* > *C. mrigala* for all TTEs and fish organs except for the Cu accumulation in the RB, where the trend was *M. armatus* > *W. attu* > *S. sarwari* > *R. rita* > *C. mrigala*. Generally, the RB of the river shows higher concentrations of all metals compared to the left bank for all species and tissues, indicating a potential difference in pollution levels or water flow patterns between the two banks. The difference is most pronounced in *W. attu*, where the increase in accumulation from the LB to RB is highly significant, especially for lead (Pb) in the liver (41.61±14.99 in LB vs. 46.03±10.56 in RB) during winter (**Table 3**).

Similar to the winter, the spring season also indicated that among all species and TTEs, the liver consistently showed the highest concentration of TTEs, followed by gills and muscles. *Wallago attu* consistently showed the highest accumulation among all metals and organs compared to other species with the significantly higher accumulation in the liver, with the value of 9.37±0.48 in LB and 13.92±0.87 in RB for Cd, 11.21±1.30 in LB and 12.36±0.47 in RB for Cu, 45.04±2.15 in LB for Pb, and 13.33±0.99 in LB and 17.92±1.46 in RB for Ni, except for the Pb (46.01±2.19) in RB which was higher in *M. armatus*. On the other hand, *C. mrigala* showed significantly lower accumulation (Cd (4.16±0.83 in RB), Cu (2.32±0.27 in LB and 3.55±0.54 in RB), Pb (11.19±0.89 in LB and 12.65±1.16 in RB), and Ni (3.59±0.40 in LB and 5.21±0.41 in RB)) in fish muscles except for the Cd concentration (5.02±0.70) in the LB which was found to be lowest in the *R. rita*. Furthermore, the pattern of TTEs’ accumulation in fish species was mostly in the order of *W. attu* > *M. armatus* > *S. sarwari* > *R. rita* > *C. mrigala* for all TTEs and fish organs in both LB and RB with the exception for the Cd accumulation in gills and muscles and Cu accumulation in liver which was in order of *W. attu* > *M. armatus* > *S. sarwari* > *C. mrigala* > *R. rita* and Ni accumulation in muscles with the order of (*M. armatus* > *W. attu* > *R. rita* >*S. sarwari* > *C. mrigala*) in the LB and gills and muscles accumulation in the Cd and Pb respectively from the RB during summer. It is observed that the RB of the river shows higher concentrations of all metals compared to the left bank for all species and tissues except for the Pb concentration in gills of *R. rita*, *W. attu*, and *M. armatus* as well as in the liver and muscles of *W. attu*. It indicates a potential difference in pollution levels or water flow patterns between the two banks. The difference is most pronounced in *M. armatus*, where the increase in accumulation from the LB to RB is highly significant, especially for lead (Pb) in the liver (38.22±3.10 in LB vs. 46.01±2.19 in RB) during spring (**Table 4**).

The accumulation TTEs during summer had a mixed trend in various organs and fish species. *Wallago attu* mostly showed the highest accumulation among all metals and organs compared to other species with the significantly higher accumulation in the liver, with the value of 23.32±0.56 in LB and 26.42±0.97 in RB for Cd, 20.72±0.42 in LB for Cu, 47.76±1.44 in LB for Pb, and 22.05±0.76 in LB and 21.94±1.55 in RB for Ni, with exception of Cu accumulation (19.99±0.35) in the RB which was higher in *S. sarwari* and Pb concentration (53.14±2.35) in the RB which was highest in *M armatus*. While, *C. mrigala* has the lowest accumulation of TTEs with the value of 5.01±1.45 in LB and 7.56±1.0 for Cd, 4.76±0.96 in LB and 5.67±0.51 in RB for Cu, 11.30±0.83 in LB and 11.80±1.47 in RB for Pb, and 4.90±0.94 in LB and 5.23±1.47 in RB for Ni in fish muscles. Moreover, the pattern of TTEs’ accumulation in studied fish was mostly in the order of *W. attu* > *M. armatus* > *S. sarwari* > *R. rita* > *C. mrigala* for most of the TTEs and fish organs in LB with a few exceptions, but it showed a mixed trend in RB during summer. It is observed that the RB of the river shows higher concentrations of all metals compared to the left bank for all species and tissues except for the Cu and Ni concentration in gills, liver, and muscles of *W. attu* and Cu and Pb concentration in the gills of *M. armatus*. The difference in concentration of TTEs between the banks is most pronounced in *C. mrigala*, where the increase in accumulation from the LB to RB is highly significant, especially for lead (Pb) in the liver (25.52±2.15 in LB vs. 37.17±3.16 in RB) during summer (**Table 5**).

**Table 3:**
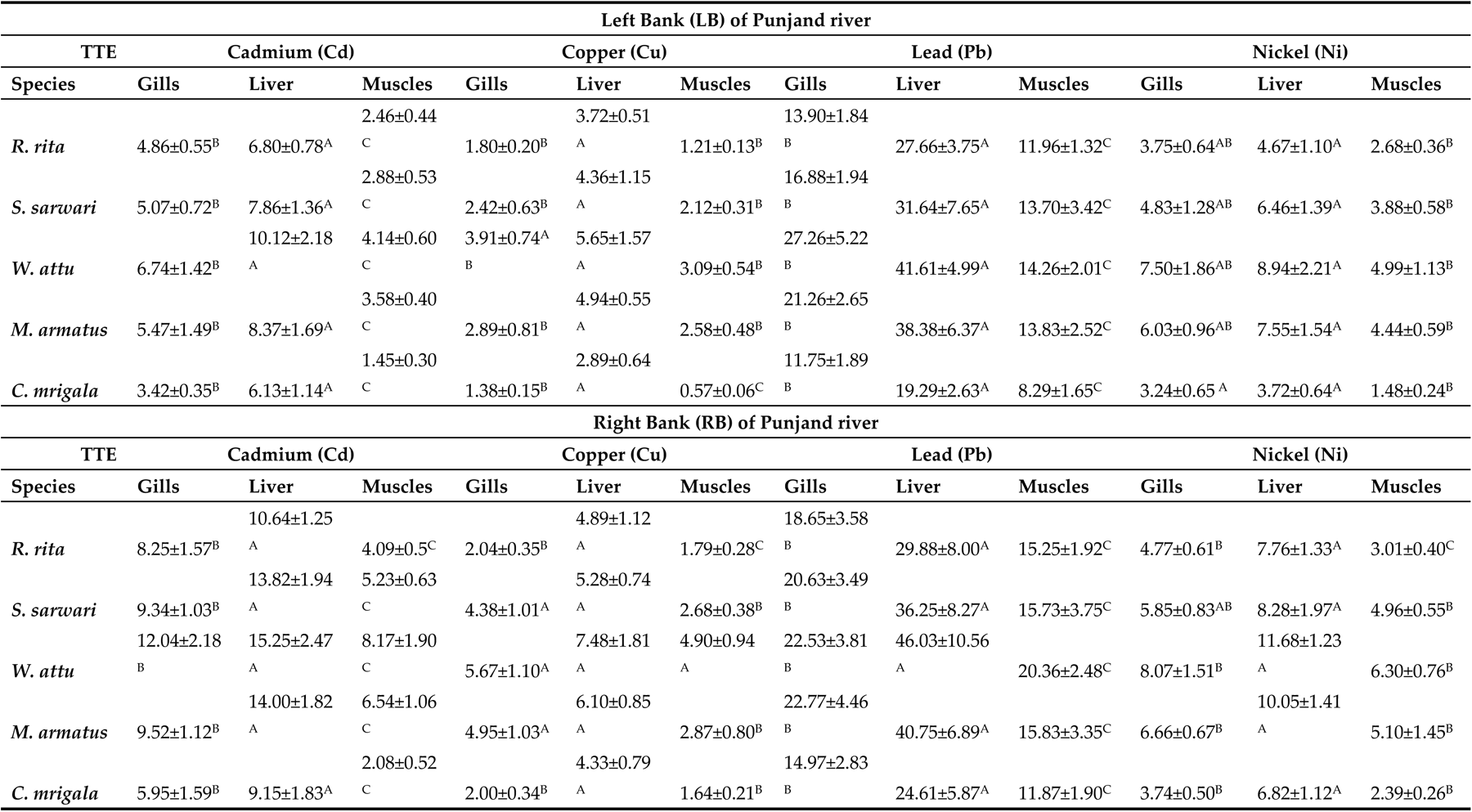
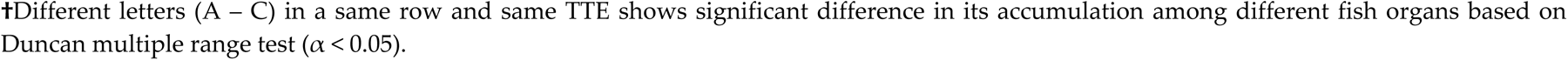
Accumulation (mg/kg) of toxic trace elements (TTEs) in different organs of five fish species collected from right (RB) and left banks (L Punjnad headworks, Uch Sharif during winter.

| Left Bank (LB) of Punjand river |  |  |  |  |  |  |  |  |  |  |  |  |
| --- | --- | --- | --- | --- | --- | --- | --- | --- | --- | --- | --- | --- |
| TTE | Cadmium (Cd) |  |  | Copper (Cu) |  |  | Lead (Pb) |  |  | Nickel (Ni) |  |  |
| Species | Gills | Liver | Muscles | Gills | Liver | Muscles | Gills | Liver | Muscles | Gills | Liver | Muscles |
| *R. rita* | 4.86±0.55<sup>B</sup> | 6.80±0.78<sup>A</sup> | 2.46±0.44<sup>C</sup> | 1.80±0.20<sup>B</sup> | 3.72±0.51<sup>A</sup> | 1.21±0.13<sup>B</sup> | 13.90±1.84<sup>B</sup> | 27.66±3.75<sup>A</sup> | 11.96±1.32<sup>C</sup> | 3.75±0.64<sup>AB</sup> | 4.67±1.10<sup>A</sup> | 2.68±0.36<sup>B</sup> |
| *S. sarwari* | 5.07±0.72<sup>B</sup> | 7.86±1.36<sup>A</sup> | 2.88±0.53<sup>C</sup> | 2.42±0.63<sup>B</sup> | 4.36±1.15<sup>A</sup> | 2.12±0.31<sup>B</sup> | 16.88±1.94<sup>B</sup> | 31.64±7.65<sup>A</sup> | 13.70±3.42<sup>C</sup> | 4.83±1.28<sup>AB</sup> | 6.46±1.39<sup>A</sup> | 3.88±0.58<sup>B</sup> |
| *W. attu* | 6.74±1.42<sup>B</sup> | 10.12±2.18<sup>A</sup> | 4.14±0.60<sup>C</sup> | 3.91±0.74<sup>A</sup> | 5.65±1.57<sup>A</sup> | 3.09±0.54<sup>B</sup> | 27.26±5.22<sup>B</sup> | 41.61±4.99<sup>A</sup> | 14.26±2.01<sup>C</sup> | 7.50±1.86<sup>AB</sup> | 8.94±2.21<sup>A</sup> | 4.99±1.13<sup>B</sup> |
| *M. armatus* | 5.47±1.49<sup>B</sup> | 8.37±1.69<sup>A</sup> | 3.58±0.40<sup>C</sup> | 2.89±0.81<sup>B</sup> | 4.94±0.55<sup>A</sup> | 2.58±0.48<sup>B</sup> | 21.26±2.65<sup>B</sup> | 38.38±6.37<sup>A</sup> | 13.83±2.52<sup>C</sup> | 6.03±0.96<sup>AB</sup> | 7.55±1.54<sup>A</sup> | 4.44±0.59<sup>B</sup> |
| *C. mrigala* | 3.42±0.35<sup>B</sup> | 6.13±1.14<sup>A</sup> | 1.45±0.30<sup>C</sup> | 1.38±0.15<sup>B</sup> | 2.89±0.64<sup>A</sup> | 0.57±0.06<sup>C</sup> | 11.75±1.89<sup>B</sup> | 19.29±2.63<sup>A</sup> | 8.29±1.65<sup>C</sup> | 3.24±0.65<sup>A</sup> | 3.72±0.64<sup>A</sup> | 1.48±0.24<sup>B</sup> |
| Right Bank (RB) of Punjand river |  |  |  |  |  |  |  |  |  |  |  |  |
| TTE | Cadmium (Cd) |  |  | Copper (Cu) |  |  | Lead (Pb) |  |  | Nickel (Ni) |  |  |
| Species | Gills | Liver | Muscles | Gills | Liver | Muscles | Gills | Liver | Muscles | Gills | Liver | Muscles |
| *R. rita* | 8.25±1.57<sup>B</sup> | 10.64±1.25<sup>A</sup> | 4.09±0.5<sup>C</sup> | 2.04±0.35<sup>B</sup> | 4.89±1.12<sup>A</sup> | 1.79±0.28<sup>C</sup> | 18.65±3.58<sup>B</sup> | 29.88±8.00<sup>A</sup> | 15.25±1.92<sup>C</sup> | 4.77±0.61<sup>B</sup> | 7.76±1.33<sup>A</sup> | 3.01±0.40<sup>C</sup> |
| *S. sarwari* | 9.34±1.03<sup>B</sup> | 13.82±1.94<sup>A</sup> | 5.23±0.63<sup>C</sup> | 4.38±1.01<sup>A</sup> | 5.28±0.74<sup>A</sup> | 2.68±0.38<sup>B</sup> | 20.63±3.49<sup>B</sup> | 36.25±8.27<sup>A</sup> | 15.73±3.75<sup>C</sup> | 5.85±0.83<sup>AB</sup> | 8.28±1.97<sup>A</sup> | 4.96±0.55<sup>B</sup> |
| *W. attu* | 12.04±2.18<sup>B</sup> | 15.25±2.47<sup>A</sup> | 8.17±1.90<sup>C</sup> | 5.67±1.10<sup>A</sup> | 7.48±1.81<sup>A</sup> | 4.90±0.94<sup>A</sup> | 22.53±3.81<sup>B</sup> | 46.03±10.56<sup>A</sup> | 20.36±2.48<sup>C</sup> | 8.07±1.51<sup>B</sup> | 11.68±1.23<sup>A</sup> | 6.30±0.76<sup>B</sup> |
| *M. armatus* | 9.52±1.12<sup>B</sup> | 14.00±1.82<sup>A</sup> | 6.54±1.06<sup>C</sup> | 4.95±1.03<sup>A</sup> | 6.10±0.85<sup>A</sup> | 2.87±0.80<sup>B</sup> | 22.77±4.46<sup>B</sup> | 40.75±6.89<sup>A</sup> | 15.83±3.35<sup>C</sup> | 6.66±0.67<sup>B</sup> | 10.05±1.41<sup>A</sup> | 5.10±1.45<sup>B</sup> |
| *C. mrigala* | 5.95±1.59<sup>B</sup> | 9.15±1.83<sup>A</sup> | 2.08±0.52<sup>C</sup> | 2.00±0.34<sup>B</sup> | 4.33±0.79<sup>A</sup> | 1.64±0.21<sup>B</sup> | 14.97±2.83<sup>B</sup> | 24.61±5.87<sup>A</sup> | 11.87±1.90<sup>C</sup> | 3.74±0.50<sup>B</sup> | 6.82±1.12<sup>A</sup> | 2.39±0.26<sup>B</sup> |

†Different letters (A – C) in a same row and same TTE shows significant difference in its accumulation among different fish organs based on Duncan multiple range test ( $\alpha < 0.05$ ).

**Table 4:** Accumulation (mg/kg) of toxic trace elements (TTEs) in different organs of five fish species collected from right (RB) and left banks of Punjnad headworks, Uch Sharif during spring.

| Left Bank (LB) of Punjand river |  |  |  |  |  |  |  |  |  |  |  |  |
| --- | --- | --- | --- | --- | --- | --- | --- | --- | --- | --- | --- | --- |
| TTE | Cadmium (Cd) |  |  | Copper (Cu) |  |  | Lead (Pb) |  |  | Nickel (Ni) |  |  |
| Species | Gills | Liver | Muscles | Gills | Liver | Muscles | Gills | Liver | Muscles | Gills | Liver | Muscles |
| *R.rita* | 6.80±0.65<sup>B</sup> | 10.79±0.81<br>A | 5.02±0.70<sup>C</sup> | 4.75±0.30<sup>B</sup> | 6.52±0.70<sup>A</sup> | 3.50±0.63<br>C | 23.44±2.42<br>B | 32.29±3.20<br>A | 14.06±2.69<sup>C</sup> | 6.62±0.69<sup>B</sup> | 9.89±0.60<sup>A</sup> | 5.24±0.52<sup>C</sup> |
| *S. sarwari* | 7.95±0.54<sup>B</sup> | 12.31±0.94<br>A | 5.78±0.62<sup>C</sup> | 6.50±1.34<sup>B</sup> | 8.78±0.75<sup>A</sup> | 4.26±0.58<br>C | 22.94±1.43<br>B | 35.10±1.89<br>A | 15.54±3.02<sup>C</sup> | 6.94±0.55<sup>B</sup> | 10.26±1.53<br>A | 5.14±0.80<sup>B</sup> |
| *W. attu* | 9.37±0.48<sup>B</sup> | 14.65±1.25<br>A | 6.65±0.53<sup>C</sup> | 8.82±0.74<sup>B</sup> | 11.21±1.30<sup>A</sup> | 6.22±0.56<br>C | 33.56±1.71<br>B | 45.04±2.15<br>A | 18.78±4.70<sup>C</sup> | 9.12±0.33<sup>B</sup> | 13.33±0.99<br>A | 6.65±0.66<sup>C</sup> |
| *M. armatus* | 8.43±1.01<sup>B</sup> | 13.60±1.83<br>A | 6.15±0.82<sup>B</sup> | 7.54±0.91<sup>B</sup> | 10.74±0.50<sup>A</sup> | 4.86±0.71<br>C | 25.28±2.10<br>B | 38.22±3.10<br>A | 15.72±0.93<sup>C</sup> | 8.50±1.04<sup>B</sup> | 12.09±0.57<br>A | 6.73±1.06<sup>B</sup> |
| *C. mrigala* | 7.07±0.31<sup>B</sup> | 10.05±0.83<br>A | 5.31±0.37<sup>C</sup> | 4.51±0.60<sup>B</sup> | 6.82±0.41<sup>A</sup> | 2.32±0.27<br>C | 14.71±1.43<br>B | 24.42±1.81<br>A | 11.19±0.89<sup>C</sup> | 6.27±0.62<sup>B</sup> | 8.32±0.95<sup>A</sup> | 3.59±0.40<sup>C</sup> |
| Right Bank (RB) of Punjand river |  |  |  |  |  |  |  |  |  |  |  |  |
| TTE | Cadmium (Cd) |  |  | Copper (Cu) |  |  | Lead (Pb) |  |  | Nickel (Ni) |  |  |
| Species | Gills | Liver | Muscles | Gills | Liver | Muscles | Gills | Liver | Muscles | Gills | Liver | Muscles |
| *R.rita* | 11.76±1.04<br>B | 15.54±0.98<br>A | 6.43±0.90<sup>C</sup> | 5.99±0.16<sup>B</sup> | 8.75±0.73<sup>A</sup> | 3.98±0.65<br>C | 20.80±1.60<br>B | 36.89±3.44<br>A | 16.12±1.15<sup>C</sup> | 8.41±1.18<sup>B</sup> | 12.02±1.05<br>A | 6.10±0.81<sup>C</sup> |
| *S. sarwari* | 13.54±0.79<br>B | 18.03±1.42<br>A | 7.42±0.97<sup>C</sup> | 8.01±0.74<sup>B</sup> | 9.58±0.51<sup>A</sup> | 5.31±0.51<br>C | 26.89±1.29<br>B | 37.61±1.78<br>A | 17.91±1.23<sup>C</sup> | 9.79±0.32<sup>B</sup> | 13.05±1.10<br>A | 6.74±0.70<sup>C</sup> |
| *W. attu* | 13.92±0.87<br>B | 20.59±2.10<br>A | 10.06±1.25<br>C | 10.16±0.64<br>B | 12.36±0.47<sup>A</sup> | 7.09±0.65<br>C | 23.81±1.38<br>B | 42.46±2.66<br>A | 17.50±1.09<sup>C</sup> | 13.07±0.51<br>B | 17.92±1.46<br>A | 7.06±0.43<sup>C</sup> |
| *M. armatus* | 12.73±0.74<br>B | 19.78±0.97<br>A | 7.86±0.65<sup>C</sup> | 8.97±0.56<sup>B</sup> | 11.01±0.69<sup>A</sup> | 5.92±0.48<br>C | 22.57±0.83<br>B | 46.01±2.19<br>A | 17.83±1.29<sup>C</sup> | 10.87±0.58<br>B | 15.33±1.07<br>A | 6.90±0.85<sup>C</sup> |
| *C. mrigala* | 8.99±0.57<sup>B</sup> | 12.26±0.74<br>A | 4.16±0.83<sup>C</sup> | 5.90±0.74<sup>B</sup> | 8.74±0.46<sup>A</sup> | 3.55±0.54<br>C | 18.66±1.94<br>B | 29.69±3.40<br>A | 12.65±1.16<sup>C</sup> | 7.92±0.45<sup>B</sup> | 11.87±0.96<br>A | 5.21±0.41<sup>C</sup> |

†Different letters (A – C) in a same row and same TTE shows significant difference in its accumulation among different fish organs based on Duncan multiple range test ( $\alpha < 0.05$ ).

**Table 5:** Accumulation (mg/kg) of toxic trace elements (TTEs) in different organs of five fish species collected from right (RB) and left banks of Punjnad headworks, Uch Sharif during summer.

| Left Bank (LB) of Punjand river |  |  |  |  |  |  |  |  |  |  |  |  |
| --- | --- | --- | --- | --- | --- | --- | --- | --- | --- | --- | --- | --- |
| TTE | Cadmium (Cd) |  |  | Copper (Cu) |  |  | Lead (Pb) |  |  | Nickel (Ni) |  |  |
| Species | Gills | Liver | Muscles | Gills | Liver | Muscles | Gills | Liver | Muscles | Gills | Liver | Muscles |
| *R. rita* | 11.43±0.55<sup>B</sup> | 15.04±0.48<sup>A</sup> | 7.47±1.10<sup>C</sup> | 7.77±0.67<sup>B</sup> | 11.63±1.00<sup>A</sup> | 5.98±0.45<sup>C</sup> | 23.60±1.25<sup>B</sup> | 36.44±1.35<sup>A</sup> | 16.88±1.72<sup>C</sup> | 10.76±0.91<sup>B</sup> | 12.80±0.38<sup>A</sup> | 7.74±0.75<sup>C</sup> |
| *S. sarwari* | 14.58±2.34<sup>A</sup> | 14.80±3.25<sup>A</sup> | 10.15±0.82<sup>A</sup> | 12.22±1.01<sup>B</sup> | 15.32±0.90<sup>A</sup> | 9.29±1.27<sup>B</sup> | 26.18±0.27<sup>B</sup> | 36.87±1.11<sup>A</sup> | 20.91±6.52<sup>C</sup> | 11.88±0.99<sup>B</sup> | 16.56±1.03<sup>A</sup> | 10.41±1.06<sup>C</sup> |
| *W. attu* | 17.46±1.87<sup>B</sup> | 23.32±0.56<sup>A</sup> | 12.96±1.18<sup>C</sup> | 16.69±2.01<sup>B</sup> | 20.72±0.42<sup>A</sup> | 14.33±0.38<sup>C</sup> | 32.60±1.78<sup>B</sup> | 47.76±1.44<sup>A</sup> | 22.99±2.64<sup>C</sup> | 17.58±1.25<sup>B</sup> | 22.05±0.76<sup>A</sup> | 15.45±0.12<sup>C</sup> |
| *M. armatus* | 14.94±0.21<sup>B</sup> | 20.58±0.90<sup>A</sup> | 12.36±1.35<sup>C</sup> | 12.90±1.20<sup>B</sup> | 18.22±0.54<sup>A</sup> | 10.49±0.34<sup>C</sup> | 28.94±1.17<sup>B</sup> | 46.60±1.52<sup>A</sup> | 21.80±1.43<sup>C</sup> | 12.71±0.36<sup>B</sup> | 17.36±0.91<sup>A</sup> | 10.69±1.01<sup>C</sup> |
| *C. mrigala* | 8.10±1.00<sup>B</sup> | 13.61±1.37<sup>A</sup> | 5.01±1.45<sup>C</sup> | 8.09±0.37<sup>B</sup> | 10.67±0.76<sup>A</sup> | 4.76±0.96<sup>C</sup> | 15.91±1.54<sup>B</sup> | 25.52±2.15<sup>A</sup> | 11.30±0.83<sup>C</sup> | 10.00±1.62<sup>B</sup> | 14.17±1.19<sup>A</sup> | 4.90±0.94<sup>C</sup> |
| Right Bank (RB) of Punjand river |  |  |  |  |  |  |  |  |  |  |  |  |
| TTE | Cadmium (Cd) |  |  | Copper (Cu) |  |  | Lead (Pb) |  |  | Nickel (Ni) |  |  |
| Species | Gills | Liver | Muscles | Gills | Liver | Muscles | Gills | Liver | Muscles | Gills | Liver | Muscles |
| *R. rita* | 15.65±1.38<sup>B</sup> | 19.91±1.36<sup>A</sup> | 10.73±0.79<sup>C</sup> | 12.72±1.51<sup>B</sup> | 15.42±0.72<sup>A</sup> | 7.7±0.48<sup>C</sup> | 27.29±0.92<sup>B</sup> | 37.09±2.19<sup>A</sup> | 19.37±0.62<sup>C</sup> | 12.52±1.38<sup>B</sup> | 17.63±0.76<sup>A</sup> | 9.20±1.47<sup>C</sup> |
| *S. sarwari* | 17.19±0.32<sup>B</sup> | 23.79±0.30<sup>A</sup> | 13.43±1.70<sup>C</sup> | 15.42±1.47<sup>B</sup> | 19.99±0.35<sup>A</sup> | 10.87±0.37<sup>C</sup> | 30.74±1.81<sup>B</sup> | 45.67±1.03<sup>A</sup> | 24.17±1.29<sup>C</sup> | 14.33±1.46<sup>B</sup> | 19.76±1.67<sup>A</sup> | 12.09±0.57<sup>C</sup> |
| *W. attu* | 22.40±1.22<sup>B</sup> | 26.42±0.97<sup>A</sup> | 13.96±1.23<sup>C</sup> | 13.48±1.22<sup>B</sup> | 17.72±0.45<sup>A</sup> | 10.79±0.92<sup>C</sup> | 34.65±1.16<sup>B</sup> | 52.02±2.31<sup>A</sup> | 29.94±7.37<sup>B</sup> | 17.39±1.14<sup>B</sup> | 21.94±1.55<sup>A</sup> | 14.20±1.28<sup>C</sup> |
| *M. armatus* | 17.33±1.14<sup>B</sup> | 22.14±1.33<sup>A</sup> | 13.39±0.91<sup>C</sup> | 14.97±0.44<sup>B</sup> | 18.07±0.56<sup>A</sup> | 11.34±1.19<sup>C</sup> | 28.30±0.97<sup>B</sup> | 53.14±2.35<sup>A</sup> | 22.67±2.61<sup>C</sup> | 18.64±0.65<sup>B</sup> | 21.53±0.88<sup>A</sup> | 12.75±0.84<sup>C</sup> |
| *C. mrigala* | 12.16±0.85<sup>B</sup> | 16.46±1.50<sup>A</sup> | 7.56±1.01<sup>C</sup> | 8.64±0.38<sup>B</sup> | 12.81±1.25<sup>A</sup> | 5.67±0.51<sup>C</sup> | 23.76±0.73<sup>B</sup> | 37.17±3.16<sup>A</sup> | 11.80±1.47<sup>C</sup> | 9.32±0.87<sup>B</sup> | 12.88±1.15<sup>A</sup> | 5.23±1.47<sup>C</sup> |

†Different letters (A – C) in a same row and same TTE shows significant difference in its accumulation among different fish organs based on Duncan multiple range test ( $\alpha < 0.05$ )

**Table 6:**
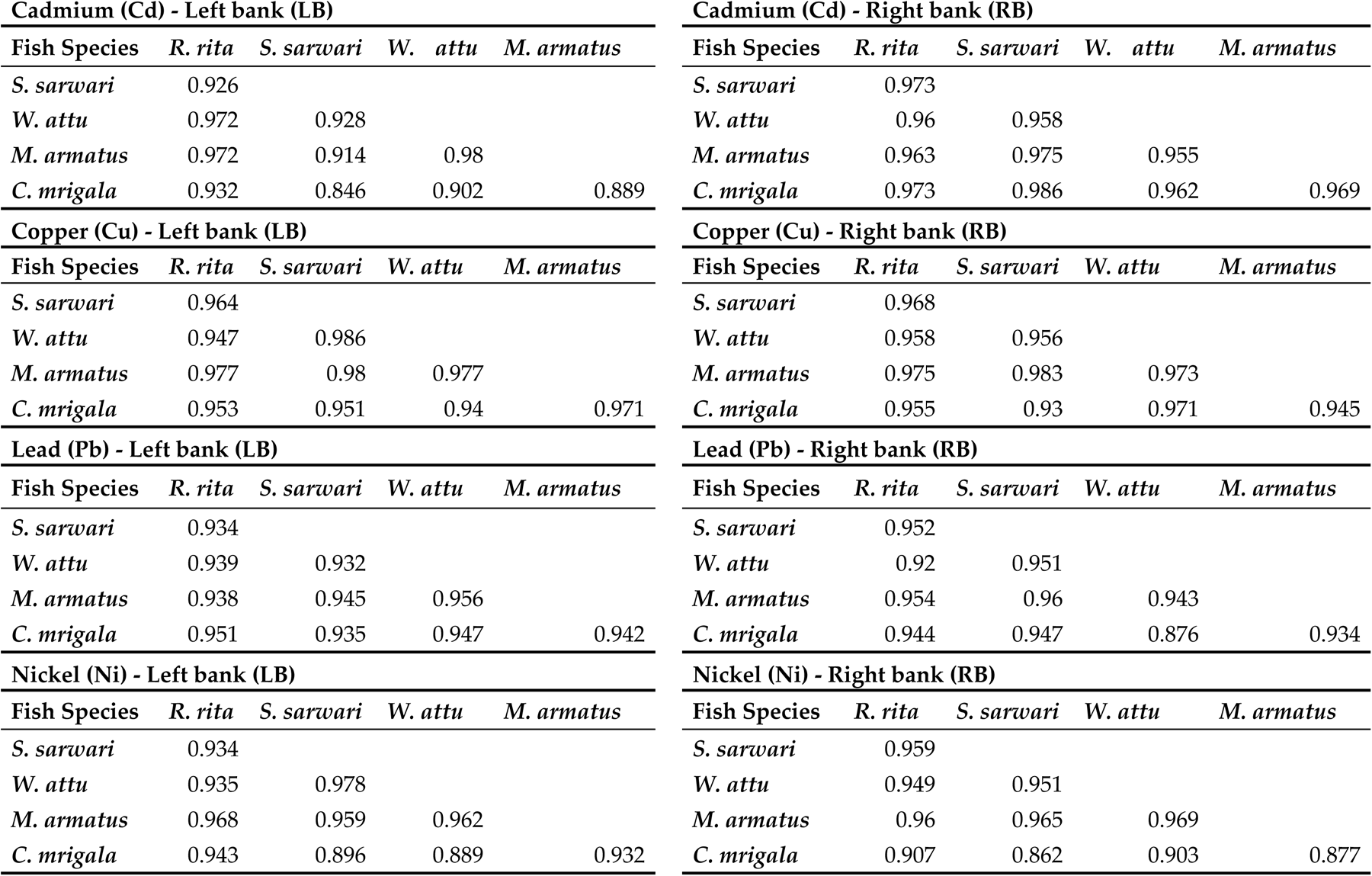
Correlation table of TTEs’ accumulation in studied fish species at left (LB) and right banks (RB) of Punjand headworks, Uch Sharif.

In order to explore the relationship between fish species and accumulation of TTEs from LB and RB, a correlation analysis was carried out. The results showed that across all metals and banks, the correlation coefficients between different fish species are generally high (above 0.90 in most cases), indicating strong positive relationships in metal accumulation. This suggests that if one species has high metal accumulation, the others are likely to show similarly high levels. The correlations are consistently high across both banks, but the right bank (RB) shows slightly higher correlations in some cases, particularly for Cadmium (Cd) and Copper (Cu). This may indicate more uniform environmental conditions on the right bank that affect all species similarly. It is more evident for the correlation for Cadmium (Cd) between *S. sarwari* and *C. mrigala* which increased from 0.846 on the left bank to 0.986 on the right bank, showing a stronger relationship on the right bank. *Wallago attu* generally exhibited the highest correlations with other species, especially for Nickel (Ni), where it shows a highly significant correlation (0.978) with ***S. sarwari*** on the left bank and strong correlations with other species on both banks. While, *C. mrigala* showed slightly lower correlations compared to other species, particularly on the left bank, which might indicate a different bioaccumulation pattern or ecological niche (**Table 6**). The correlation analysis reveals that the accumulation of toxic trace elements in fish species from the Punjand headworks is highly interrelated, with particularly strong correlations observed on the right bank. This suggests that environmental factors influencing metal accumulation are relatively uniform across species, especially on the right bank. The consistent high correlations underline the importance of monitoring multiple species to understand the broader environmental impact of heavy metal contamination in this area.

The difference in the concentration of TTEs between the LB and RB of the river was visualized using box plot. The results indicated that among all species, lead (Pb) showed the highest accumulation compared to other toxic trace elements (Cd, Cu, Ni), especially in the RB samples. This is evident from the box plots where Pb’s range and median values significantly exceed those of the other elements or most species. Moreover, the RB of the Punjand River generally exhibits higher bioaccumulation levels across all metals. This trend is particularly noticeable in *W. attu* and *S. sarwari*, where the difference is marked, indicating a higher contamination level on the right bank. Although variations exist, the general pattern of higher accumulation on the right bank is consistent across species, reinforcing the environmental impact on this side of the river. Among various fish species, *W. attu* showed the most significant accumulation for all elements, particularly for Pb and Ni, indicating that this species is highly susceptible to heavy metal bioaccumulation while, *C. mrigala* and *R. rita* displayed lower bioaccumulation levels, suggesting either a different habitat preference or varying metabolic rates in processing these elements (**Figure 2**). The box plots analysis illustrated a clear trend of higher heavy metal bioaccumulation in fish species from the right bank of the Punjand River, with Pb being the most prominent contaminant.

**Figure 2.**
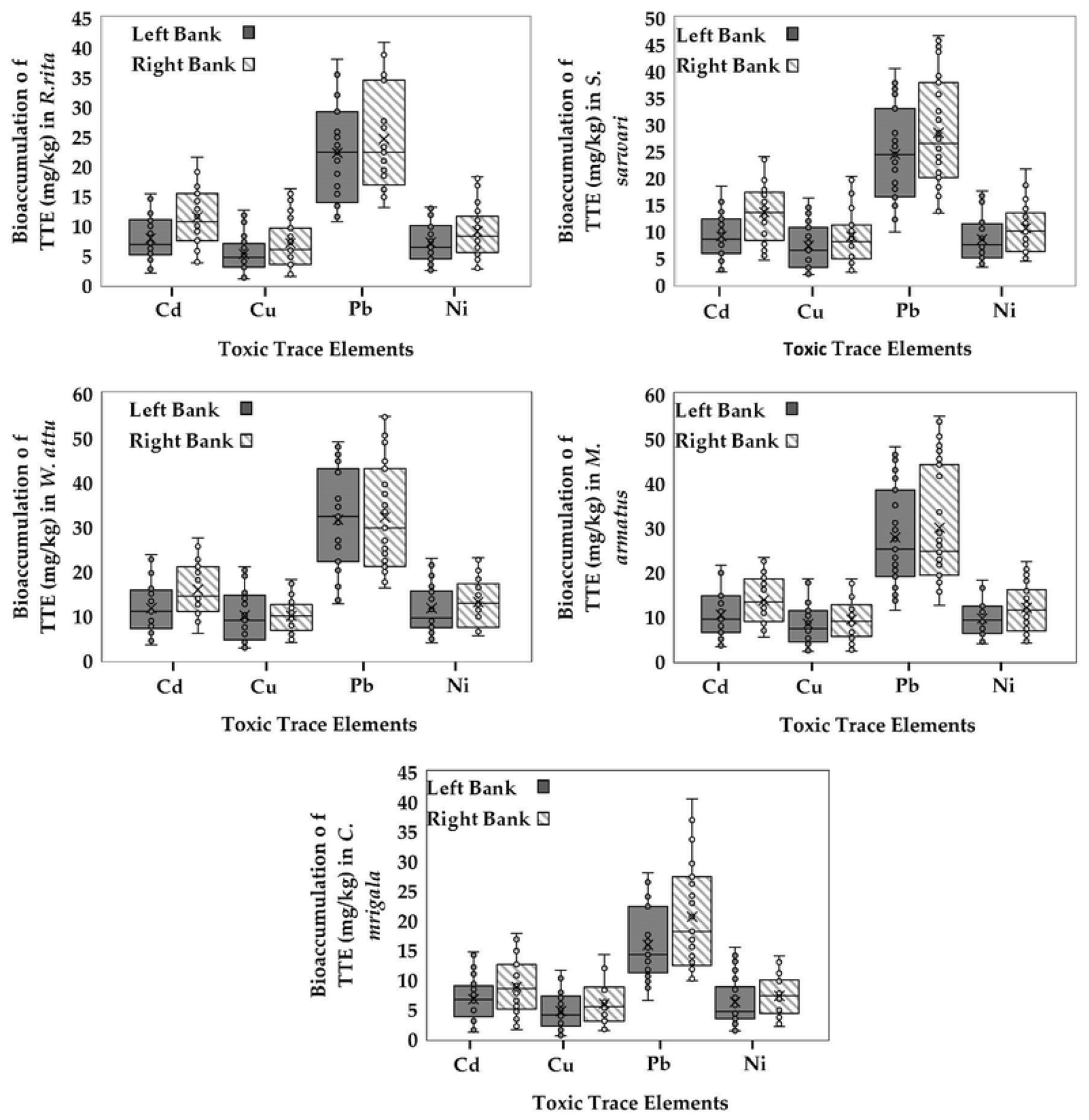
Comparison of accumulation of four toxic trace elements (TTEs) (Cd, Cu, Pb, and Ni) in five fish species Between Left and Right Banks of Punjand headworks

The season wise comparison of TTEs’ accumulation in different fish at LB of Punjand headworks showed a significant difference in accumulation of TTEs across three seasons. The results showed that during summer maximum accumulation of all trace elements in all species was observed. *W. attu* showed highest accumulation of Pb during summer while *C. mrigala* showed the lowest concentration of Cu during winter. Overall, the trend of accumulation of TTEs increased gradually from winter to summer indicating a direct relationship of accumulation of TTEs and temperature of aquatic ecosystem (**Figure 3-A**). Similar to the LB, RB also presented an increasing trend of TTEs accumulation from winter to summer. The maximum accumulation of TTEs was observed in *W. attu*, with the maximum concentration of Pb during summer. Winter season showed overall low concentration of all TTEs across all studies fish. The lowest concentration of TTEs was determined in *C. mrigala* during winter for Cu concentration. Spring usually showed moderate accumulation of TTEs in all fish species (**Figure 3-B**).

**Figure 3-A.**
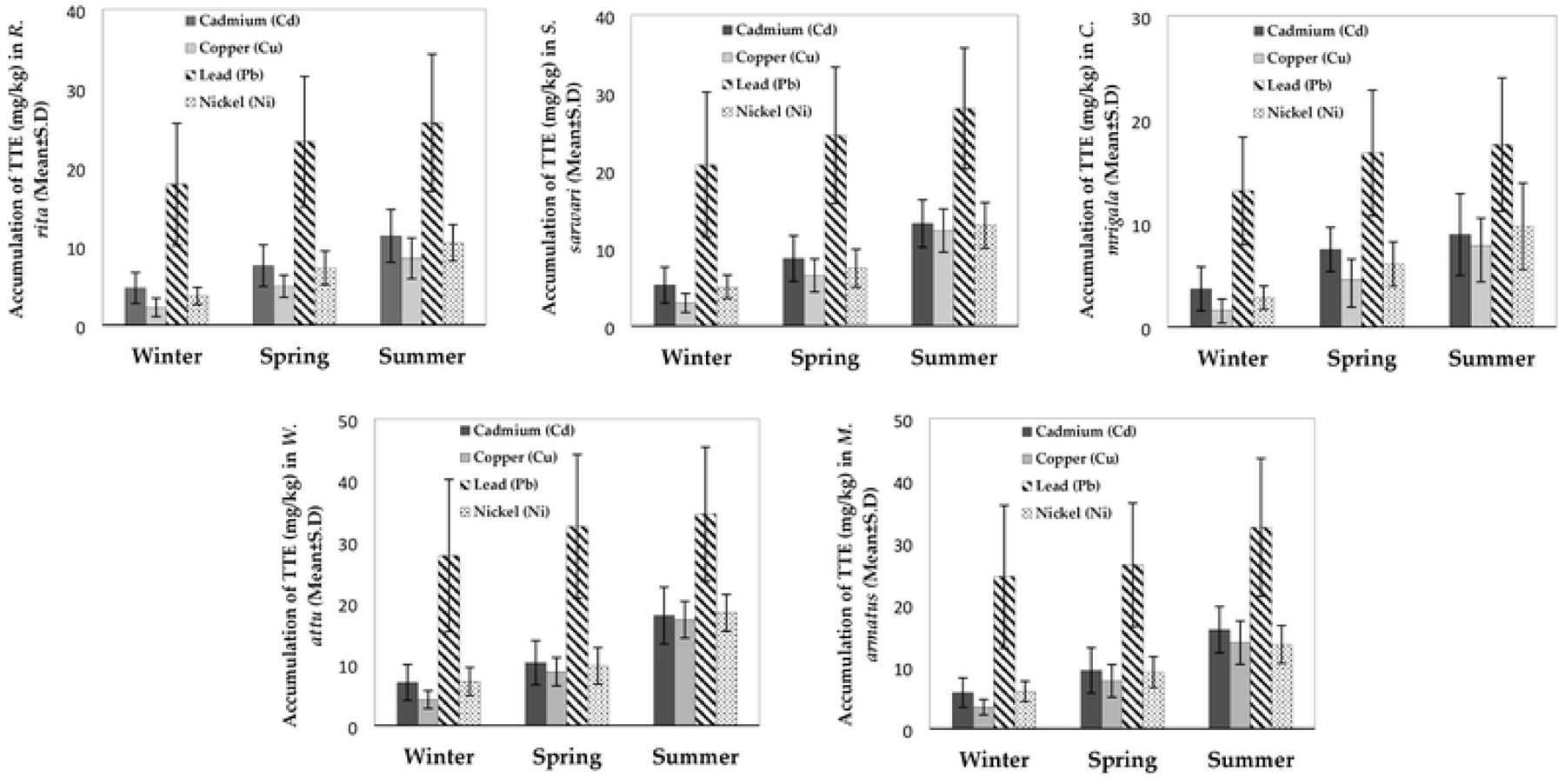
Accumulation comparison of four toxic trace elements (TTEs) (Cd, Cu, Ni, and PB) in five fish species (*R. rita*, *S. sarwari*, *W. attu*, *M. armatus*, and *C. mrigala*) between three different sampling seasons (winter, spring, and summer) at the left bank (LB) of Punjnad headworks.

**Figure 3-B:**
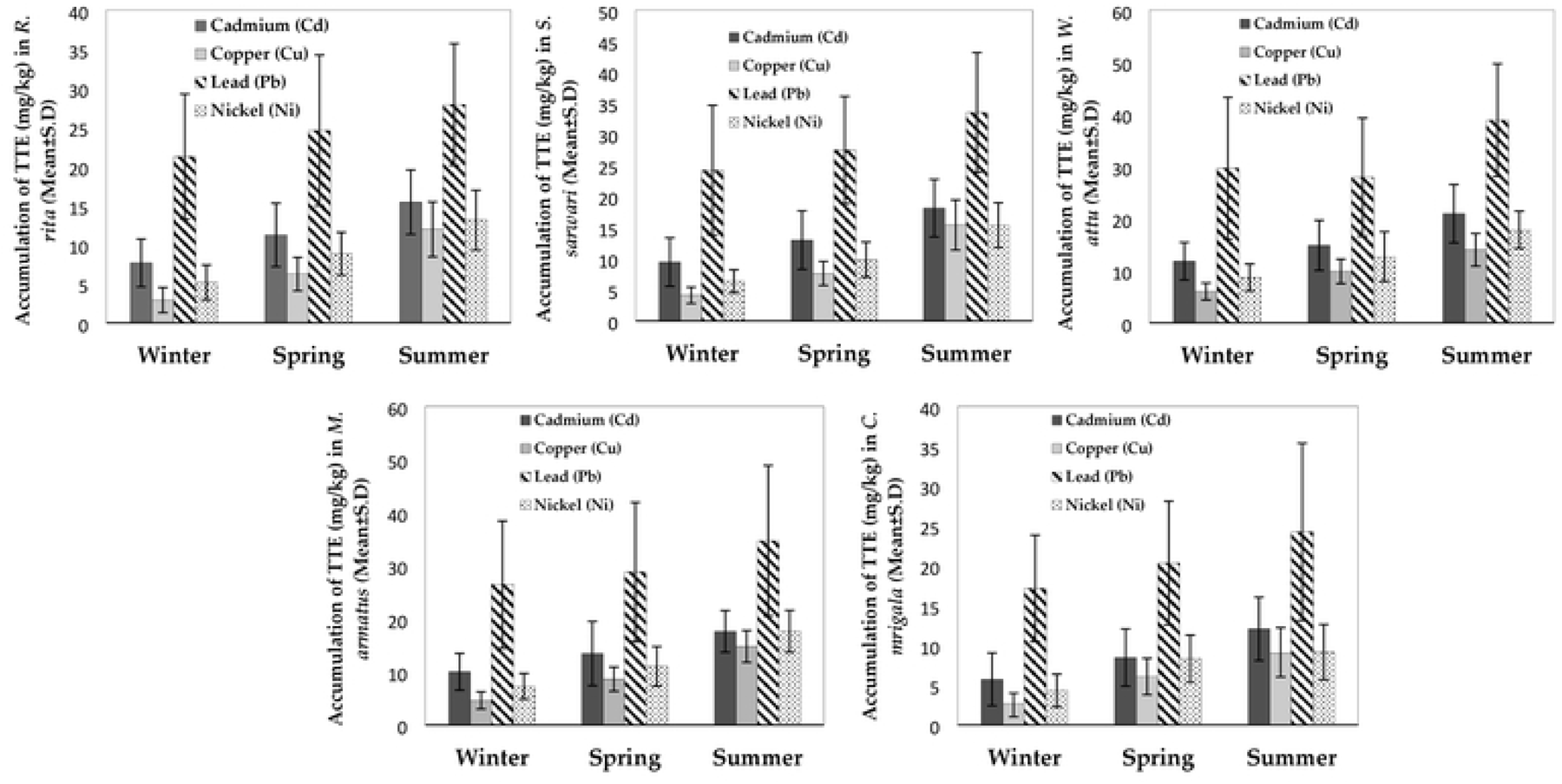
Accumulation comparison of four toxic trace elements (TTEs) (Cd, Cu, Ni, and PB) in five fish species (*R. rita*, *S. sarwari*, *W. attu*, *M. armatus*, and *C*. *mrigala*) between three different sampling seasons (winter, spring, and summer) at the right bank (RB) of Punjnad headworks

Current study applied the Hierarchical cluster analysis (HCA) with the Wards linkage method and Euclidean distance as parameters to measure the similarity/variation for accumulation of TTEs in fish. HCA of fish species and the TTEs formed three clusters (group - 1, group – 2, group – 3). Among fish species, cluster one (group - 1) is formed due to the high content of TTEs in *W. attu* – LB, *W. attu* - RB, *S, sarwari* – RB, and *M. armatus* - RB. Cluster two (group - 2) explained the moderate concentration of TTEs in *M. armatus* – LB, *S. sarwari* – LB, and *R. rita* – RB during the sampling seasons. Cluster three (group - 3) is formed to indicate the lowest accumulation of TTEs in *R. rita* – LB, *C. mrigala* – RB, and *C. mrigala* – LB (**Figure 4-A**). For the TTEs, Cluster 1 (group – 1) showed the lowest concentration of Cd – LB, Ni – LB, Cu – LB and Cu – RB in fish at the Punjnad river. Cluster 2 (group – 2) is formed due to the moderate accumulation of Ni – RB and Cd – Rb at the study site. The formation of three different groups showed that Pb concentration was significantly higher at LB and RB of the river forming group – 3 (Figure 4-B).

**Figure 4:**
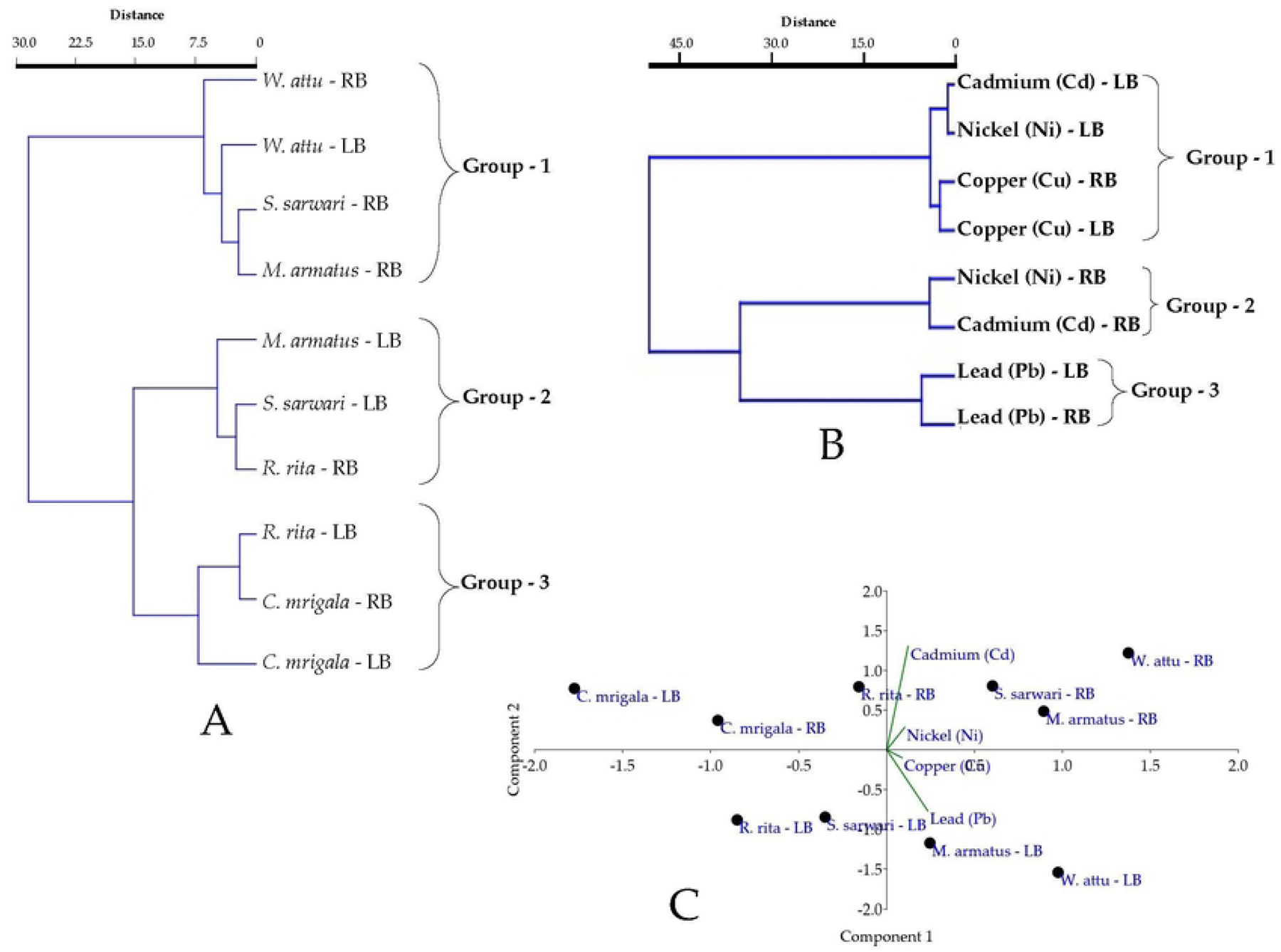
Principal Component Analysis of Accumulation of Toxic Trace Elements (TTEs) (Cd, Cu, Pb, and Ni) in Five Fish Species (*R. rita*, *S. sarwari*, *W. attu*, *M. armatus*, and *C. mrigala*) from Left (LB) and Right Banks (RB) of Punjnad Headworks.

The study also carried out the principal component analysis (PCA) to demonstrate the similarity/variation in distribution and behavior of TTEs and studied fish species. The obtained results showed, two principal components (PCs) with Eigenvalue > 1 and two PC having Eigenvalue < 1. The Eigenvalues value of PC1, PC2, PC3, and PC4 were found to be 42.85, 1.57, 0.24, and 0.08 with the variance of 95.75, 3.51, 0.54, and 0.20%, respectively. It is observed that PC1 was positively dominated by Pb having loading value of 0.79, Cd dominated the PC2 having loading value of 0.85, PC3 was dominated by Cu having a loading value of 0.73, a negatively dominated Ni with loading value of -0.76 was indicated for PC4 (**Figure 4C**).

### 3.2 Human Health Risk Assessment of TTEs

Human health risk assessment of toxic trace elements (TTEs) in fish muscles is important to highlight the bioaccumulation and harmful effects of trace elements like Cd, Cu, Pb, and Ni, which can pose serious health risks to consumers. Understanding the levels of TTEs in fish helps in identifying potential hazards, guiding regulatory limits, and ensuring food safety, thereby protecting public health from long-term exposure to toxic substances. Current study evaluated the potential hazard of TTEs present in fish muscles to the mature regular consumers (MRC), juvenile regular consumers (JRC), mature seasonal consumers (MSC) and juvenile seasonal consumers (JSC). Overall, the results showed that the potential hazard of TTEs was in the order of JRC > MRC > JSC > MSC in all fish species and studied TTEs. The maximum exposure hazard was found to the JRC in *W. attu* for Pb accumulation (0.071) while the minimum exposure hazard was recorded to the MSC in *C. mrigala* for Ni concentration (0.006) in fish muscles.

The trend of potential exposure hazard of TTEs was in the order of Pb > Ni > Cd > Cu for *R. rita*, *S. sarwari*, and *W. attu*. While, in *M. armatus* and *C mrigala* the trend was in order of Pb > Cd > Cu > Ni. Across species, the exposure hazard of Cd and Ni was in order of *M. armatus* > *W. attu* > *S. sarwari* > *R. rita* > *C. mrigala*, the potential exposure hazard of Pb was in the order *W. attu* > *M. armatus* > *S. sarwari* > *R. rita* > *C. mrigala*, and for Ni the trend was in the order of *W. attu* > *S. sarwari* > *M. armatus* > *R. rita* > *C. mrigala*.

The analysis of THQ showed that for all fish species, Cd had the highest THQ values across all consumer groups, and it was in the order of *M. armatus* > *W. attu* > *S. sarwari* > *R. rita* > *C. mrigala*. It possessed more health risks for regular consumers (both adults and juveniles), indicating a significant health risk due to Cd exposure. The THQ values for copper are relatively low across all species and consumer groups, with the order of *M. armatus* > *W. attu* > *S. sarwari* > *R. rita* > *C. mrigala* suggesting a lower health risk associated with copper exposure. Lead (Pb) also poses a considerable health risk, particularly to juvenile regular consumers, although its THQ values are generally lower than those for Cd. Its exposure across species was in the order of *W. attu* > *M. armatus* > *S. sarwari* > *R. rita* > *C. mrigala*. The THQ values for Ni the lowest with a trend of *W. attu* > *S. sarwari* > *M. armatus* > *R. rita* > *C. mrigala*. Overall, the highest THQ was observed in Cd (42.826) in the muscles of *M. armatus* with the potential hazard to the JRC. The lowest THQ effects were observed in *R. rita* muscles affecting MSC in Cu (0.190). The total hazard quotient (TTHQ) values indicate that regular consumers, especially juveniles, are at a higher risk compared to seasonal consumers. Among the fish species listed, *W. attu* and *M. armatus* show the highest TTHQ values (59.56 and 55.33 respectively), signaling that these species are the most hazardous to consume regularly followed by *S. sarwari*, *R. rita*, and *C. mrigala* with TTHQ values of 43.3, 40.41, and 34.54 respectively. Metal Pollution Index of TTEs (MPI_TTE_) highlighted the overall contamination level found in *M armatus* and *W. attu* (11.88 and 11.71 respectively showing significantly more contamination of TTEs, followed by *S. sarwari*, *R. rita*, and *C. mrigala* with the MPI values of 9.56, 8.81, and 7.1 respectively. show higher MPI values, indicating higher levels of metal contamination (**Table 7**). The assessment indicates that consuming these fish species, particularly on a regular basis, poses a health risk due to the presence of toxic trace elements, especially cadmium and lead. Regular consumers, particularly juveniles, are more at risk than seasonal consumers.

## 4. Discussion

Current study explored the accumulation of selected TTEs (Cd, Cu, Pb, and Ni) in three organs (liver, gills, and muscles) of five fish species (*R. rita*, *S. sarwari*, *W. attu*, *M. armatus*, and *C. mrigala* , during winter, spring and summer collected from LB and RB of Punjnad headworks. Furthermore, the health risk assessment of these TTEs to human health was also determined to explore its toxic effects on mature and juvenile consumers. A significantly, lower accumulation of Cu was indicated in this study. In accordance to our findings, a study was conducted to measure Cu and Zn concentrations in fish samples from three rivers in the Malakand Division, Pakistan. The muscles of *Mastacembelus armatus* from Chakdara on the river Swat exhibited the highest Cu concentration. The study indicated that Zn concentrations were generally higher than Cu in fish organs (44). Our results showed that *M. armatus* has higher affinity to accumulate TTEs. Similar to our findings, a study was conducted to determine the TTEs in three fish species *M. armatus, Channa punctatus, and Glossogobius giuris* from the Turag River in Bangladesh. The study measured concentrations of Pb, Cd, chromium (Cr), Cu, and iron (Fe) in the fish and found higher concentration of TTEs (Cd and Pb) in *M. armatus* (45). Lower concentrations of Cu and Ni in our study is coherent with a study that investigated the bioaccumulation of heavy metals viz., Fe, Ni, manganese (Mn), zinc (Zn), and Cu in various tissues of *M. armatus*, including the gills, liver, kidneys, muscles, and integument, from a rivulet in Kasimpur and concluded lower levels of Cu and Ni accumulation in various organs of the fish (46). Out study showed a high level of Pb accumulation in fish organs which was also been reported in a study by (47), showing high level of Pb accumulation in various organs of fish collected from the Punjnad headworks. However, in contrast to our findings, a recent investigation focused on the bioaccumulation of potentially toxic elements (PTEs) including Cd, cobalt (Co), Cr, Cu, lithium (Li), Ni, Pb, selenium (Se), Zn, and Mn in 12 economically important fish species from the lower Ganges in India found that the average concentrations of these potentially toxic elements (PTEs) were ranked as: Zn > Cu > Mn > Ni > Se > Cr > Pb > Co ∼ Li > Cd (42). This could be explained by the nature of these freshwater as out study site has more stagnant water as compared to this study which could lead to a different pattern of accumulation for TTEs.

**Table 7.** Human Health Risk Assessment of Toxic Trace Elements (TTEs) present in the muscles of studied fish species. The human health risk assessment is in the order of *R. rita, S. sarwari, W. attu, M. armatus, and C. mrigala* respectively.

| TTEs | EXP<sup>r</sup><sub>M</sub> | EXP<sup>r</sup><sub>J</sub> | EXP<sup>s</sup><sub>M</sub> | EXP<sup>s</sup><sub>J</sub> | THQ<sup>r</sup><sub>M</sub> | THQ<sup>r</sup><sub>J</sub> | THQ<sup>s</sup><sub>M</sub> | THQ<sup>s</sup><sub>J</sub> | TTHQ<sup>r</sup><sub>M</sub> | TTHQ<sup>r</sup><sub>J</sub> | TTHQ<sup>s</sup><sub>M</sub> | TTHQ<sup>s</sup><sub>J</sub> | MPI<sub>TTE</sub> |
| --- | --- | --- | --- | --- | --- | --- | --- | --- | --- | --- | --- | --- | --- |
| **Cadmium (Cd)** | 0.021 | 0.025 | 0.009 | 0.010 | 21.17 | 25.194 | 8.892 | 10.374 |  |  |  |  |  |
| **Copper (Cu)** | 0.018 | 0.022 | 0.008 | 0.009 | 0.452 | 0.538 | 0.190 | 0.222 | 33.96 | 40.41 | 14.26 | 16.64 | 8.81 |
| **Lead (Pb)** | 0.045 | 0.053 | 0.019 | 0.022 | 11.162 | 13.283 | 4.688 | 5.470 |  |  |  |  |  |
| **Nickel (Ni)** | 0.023 | 0.028 | 0.010 | 0.012 | 1.175 | 1.398 | 0.493 | 0.576 |  |  |  |  |  |
| TTEs | EXP<sup>r</sup><sub>M</sub> | EXP<sup>r</sup><sub>J</sub> | EXP<sup>s</sup><sub>M</sub> | EXP<sup>s</sup><sub>J</sub> | THQ<sup>r</sup><sub>M</sub> | THQ<sup>r</sup><sub>J</sub> | THQ<sup>s</sup><sub>M</sub> | THQ<sup>s</sup><sub>J</sub> | TTHQ<sup>r</sup><sub>M</sub> | TTHQ<sup>r</sup><sub>J</sub> | TTHQ<sup>s</sup><sub>M</sub> | TTHQ<sup>s</sup><sub>J</sub> | MPI |
| **Cadmium (Cd)** | 0.023 | 0.027 | 0.009 | 0.011 | 22.596 | 26.891 | 9.491 | 11.073 |  |  |  |  |  |
| **Copper (Cu)** | 0.019 | 0.022 | 0.008 | 0.009 | 0.472 | 0.561 | 0.198 | 0.231 | 36.38 | 43.3 | 15.28 | 17.83 | 9.56 |
| **Lead (Pb)** | 0.048 | 0.057 | 0.020 | 0.023 | 11.948 | 14.219 | 5.019 | 5.855 |  |  |  |  |  |
| **Nickel (Ni)** | 0.027 | 0.033 | 0.011 | 0.013 | 1.368 | 1.628 | 0.575 | 0.670 |  |  |  |  |  |
| TTEs | EXP<sup>r</sup><sub>M</sub> | EXP<sup>r</sup><sub>J</sub> | EXP<sup>s</sup><sub>M</sub> | EXP<sup>s</sup><sub>J</sub> | THQ<sup>r</sup><sub>M</sub> | THQ<sup>r</sup><sub>J</sub> | THQ<sup>s</sup><sub>M</sub> | THQ<sup>s</sup><sub>J</sub> | TTHQ<sup>r</sup><sub>M</sub> | TTHQ<sup>r</sup><sub>J</sub> | TTHQ<sup>s</sup><sub>M</sub> | TTHQ<sup>s</sup><sub>J</sub> | MPI |
| **Cadmium (Cd)** | 0.029 | 0.035 | 0.012 | 0.014 | 29.399 | 34.986 | 12.348 | 14.406 |  |  |  |  |  |
| **Copper (Cu)** | 0.021 | 0.025 | 0.009 | 0.010 | 0.515 | 0.613 | 0.216 | 0.253 | 46.5 | 55.33 | 19.53 | 22.78 | 11.71 |
| **Lead (Pb)** | 0.059 | 0.071 | 0.025 | 0.029 | 14.839 | 17.659 | 6.233 | 7.272 |  |  |  |  |  |
| **Nickel (Ni)** | 0.035 | 0.041 | 0.015 | 0.017 | 1.743 | 2.075 | 0.732 | 0.854 |  |  |  |  |  |
| TTEs | EXP<sup>r</sup><sub>M</sub> | EXP<sup>r</sup><sub>J</sub> | EXP<sup>s</sup><sub>M</sub> | EXP<sup>s</sup><sub>J</sub> | THQ<sup>r</sup><sub>M</sub> | THQ<sup>r</sup><sub>J</sub> | THQ<sup>s</sup><sub>M</sub> | THQ<sup>s</sup><sub>J</sub> | TTHQ<sup>r</sup><sub>M</sub> | TTHQ<sup>r</sup><sub>J</sub> | TTHQ<sup>s</sup><sub>M</sub> | TTHQ<sup>s</sup><sub>J</sub> | MPI |
| **Cadmium (Cd)** | 0.036 | 0.043 | 0.015 | 0.018 | 35.986 | 42.826 | 15.115 | 17.634 |  |  |  |  |  |
| **Copper (Cu)** | 0.031 | 0.037 | 0.013 | 0.015 | 0.784 | 0.933 | 0.329 | 0.384 | 50.04 | 59.56 | 21.02 | 24.52 | 11.88 |
| **Lead (Pb)** | 0.048 | 0.057 | 0.020 | 0.024 | 12.055 | 14.347 | 5.064 | 5.907 |  |  |  |  |  |
| **Nickel (Ni)** | 0.024 | 0.029 | 0.010 | 0.012 | 1.218 | 1.450 | 0.512 | 0.597 |  |  |  |  |  |
| TTEs | EXP<sup>r</sup><sub>M</sub> | EXP<sup>r</sup><sub>J</sub> | EXP<sup>s</sup><sub>M</sub> | EXP<sup>s</sup><sub>J</sub> | THQ<sup>r</sup><sub>M</sub> | THQ<sup>r</sup><sub>J</sub> | THQ<sup>s</sup><sub>M</sub> | THQ<sup>s</sup><sub>J</sub> | TTHQ<sup>r</sup><sub>M</sub> | TTHQ<sup>r</sup><sub>J</sub> | TTHQ<sup>s</sup><sub>M</sub> | TTHQ<sup>s</sup><sub>J</sub> | MPI |
| **Cadmium (Cd)** | 0.020 | 0.023 | 0.008 | 0.010 | 19.594 | 23.318 | 8.230 | 9.602 |  |  |  |  |  |
| **Copper (Cu)** | 0.019 | 0.023 | 0.008 | 0.009 | 0.482 | 0.573 | 0.202 | 0.236 | 29.03 | 34.54 | 12.19 | 14.22 | 7.1 |
| **Lead (Pb)** | 0.033 | 0.039 | 0.014 | 0.016 | 8.275 | 9.847 | 3.476 | 4.055 |  |  |  |  |  |
| **Nickel (Ni)** | 0.014 | 0.016 | 0.006 | 0.007 | 0.676 | 0.805 | 0.284 | 0.331 |  |  |  |  |  |

†EXP<sup>r</sup><sub>M</sub> = Exposure rate in adult regular consumers, EXP<sup>r</sup><sub>J</sub> = Exposure rate in juvenile regular consumers, EXP<sup>s</sup><sub>M</sub> = Exposure rate in adult seasonal consumers, EXP<sup>s</sup><sub>J</sub> = Exposure rate in juvenile seasonal consumers, THQ<sup>r</sup><sub>M</sub> = Target hazard quotient in adult regular consumers, THQ<sup>r</sup><sub>J</sub> = Target hazard quotient in juvenile regular consumers, THQ<sup>s</sup><sub>M</sub> = Target hazard quotient in adult seasonal consumers, THQ<sup>s</sup><sub>J</sub> = Target hazard quotient in juvenile seasonal consumers, TTHQ<sup>r</sup><sub>M</sub> = Total Target hazard quotient in adult regular consumers, TTHQ<sup>r</sup><sub>J</sub> = Total Target hazard quotient in juvenile regular consumers, TTHQ<sup>s</sup><sub>M</sub> = Total Target hazard quotient in adult seasonal consumers, TTHQ<sup>s</sup><sub>J</sub> = Total Target hazard quotient in juvenile seasonal consumers MPI = Metal pollution index

In a relatable study, the bioaccumulation of heavy metals arsenic (As), mercury (Hg), and Cd was investigated in five fish species: *Ctenopharyngodon idella, Oreochromis niloticus, Eutropiichthys vacha, R. rita*, and *S. sarwari* from Head Punjnad, Pakistan. The findings indicated that the metal concentrations were ranked as follows: Cd > As > Hg. Specifically, *R. rita, O. niloticus*, and *C. idella* demonstrated elevated levels of metal accumulation, suggesting that species-specific bioaccumulation patterns may be influenced by their feeding behaviors (48). High level of Cd and Pb accumulation in fish organs has also been reported in another study which concluded that in various fish species Cd and Pb had the highest accumulation in contrast to other TTEs (49). Our study reported a higher accumulation of Pb and Cd in comparison to Ni and Cu. Similar findings have been reported in a study which explored the concentration of TTEs in various organs of three edible fish species, *Catla catla, W, attu*, and *Tilapia nilotica* and found higher accumulation of Pb and Cd than Ni and Cu (50). Ali and Khan (51) has studied the bioaccumulation of Cr, Ni, Cd, and Pb in the carnivorous fish *M. armatus* across various sites in three rivers within the Malakand Division, Pakistan. The study revealed that Pb concentrations in the muscles of *M, armatus* was found to be highest followed by Cd, which is also reported in current findings. Furthermore, the liver was found to accumulate

Fish are particularly prone to heavy metal accumulation due to their efficient absorption of metals through the gills, which are involved in both respiration and excretion. The direct exposure of the gills to water facilitates the uptake of environmental metals, highlighting the importance of monitoring heavy metal levels to assess the health risks associated with fish consumption (20). The accumulation of heavy metals in fish, particularly in critical organs such as the liver, gills, and muscles, is of significant concern due to the potential health risks posed by consuming contaminated fish. These organs tend to retain varying amounts of heavy metals, underscoring the importance of continuous monitoring of metal levels in fish. Such ongoing surveillance is crucial for raising awareness about the potential dangers associated with consuming fish contaminated with heavy metals (52).

Our study reported exceeded amount of Cd and Pb for their safe human consumption according to the safety limits recommended by the FAO/WHO, indicating a significant risk of metal exposure which is in accordance to another study that found that the liver had the highest concentration and TTEs accumulation in muscles exceeded the safety limits recommended by the FAO/WHO, indicating a significant risk of metal (46). Similarly, Pandey, Pandey (53), also showed exceeded concentrations of Cr, Cd, and Pb in the muscles of *R. rita* in comparison to the limits set by the Food and Agriculture Organization (FAO) and the U.S. Environmental Protection Agency (USEPA). In a recent study by Anwarul Hasan, Satter (54), heavy metals including Cu, Zn, Pb, Cd, Ni, and As were analyzed in three common fish species viz., *Systomus sarana, Pethia ticto, and M. armatus* from the Shitalakshya river using atomic absorption spectroscopy (AAS). The study found that concentrations of Cu, Zn, Pb, Cd, Ni, and As in these fish exceeded the international safety standards set by the FAO/WHO, the U.S. Food and Drug Administration (USFDA), the Ministry of Food and Livestock (MOFL), and the European Commission (EC). Although the targeted hazard quotient (THQ) values for these metals were within the limits deemed safe for individual exposure, the cumulative hazard index (HI) for all three fish species surpassed acceptable levels, indicating a potential health risk for consumers.

According to Naz, Chatha (47) *C. mrigala* from Punjnad headworks was found to be unsafe for human consumption due to elevated levels of the total target hazard quotient (TTHQs) which is in coherence to current findings. In contrast to our findings, a study concluded that the metal pollution index (MPI), targeted hazard quotient (THQ), and total target hazard quotient (TTHQs) TTEs were below 1, indicating minimal health risks associated with fish consumption from this area total target hazard quotient (TTHQs) (42). Similarly, in a related study, the contamination levels of major rivers, including the Ravi, Chenab, Kabul, and Indus, were assessed, with a particular focus on their impact on fish and human health. The River Ravi, heavily polluted by industrial and sewage wastewater, was identified as the most contaminated, posing significant risks to aquatic life and human health. In contrast, the Indus River, with its larger water volume and fewer industrial sources, demonstrated better ecosystem health. Fish from the Indus, Chenab, and Jhelum Rivers were deemed safe for human consumption. The study underscored the critical need for effective wastewater treatment to mitigate the harmful effects of heavy metals on both aquatic ecosystems and public health (55). This difference in acceptability of fish for human consumption could be due to geographical location and increased human activity as the current study site as compared to the other reported sites. It is important to consider this drastic change in accumulation of TTEs and the issue must be delt with urgency to reduce the accumulation of TTEs in water bodies of areas reported in the current study.

## 5. Conclusions

The findings of this study underscore the critical impact of industrial and agricultural runoff on the aquatic ecosystem at the Punjnad Headworks, where toxic trace elements (TTEs) are accumulating in significant quantities. The bioaccumulation of Cd, Cu, Pb, and Ni in the liver, gills, and muscles of fish species like *R. rita*, *S. sarwari*, *W. attu*, *M. armatus*, and *C. mrigala* was particularly concerning, with *Wallago attu* showing the highest levels of TTEs. Human health risk assessments indicated that both Cd and Pb posed the most considerable exposure hazards, with Target Hazard Quotients (THQs) exceeding safe limits, particularly for these metals. The Total Target Hazard Quotients (TTHQs) also indicated that consumption of fish from the study sites poses a considerable risk to human health. The Metal Pollution Index (MPI) suggested moderate to high levels of contamination across all fish species studied. The study concluded that the right bank of the Punjnad headworks is more heavily contaminated than the left, and fish consumption from both banks is unsafe due to the elevated levels of toxic trace elements. These results highlight an urgent need for remedial actions to mitigate TTE pollution and protect both aquatic life and human health.

## Acknowledgements

We are thankful to the Directorate of Fish Hatchery, Bahawalpur, for assisting in netting and sample collection. We are also thankful to the Mian Nawaz Sharif University of Agriculture for analyzing fish samples.

## Ethics declarations

### Ethics approval and consent to participate

This study was approved by the Department Committee on Animal Ethics and Welfare, Government Sadiq College Women University, Bahawalpur Pakistan. All the authors have equally participated in this study.

### Consent for publication

All the authors agreed to publish the work in this journal.

### Author Contributions

Conceptualization, Saima Mustafa and Ahmad Chatha; Data curation, Qudrat Ullah, Dalia Fouad, Maria Lateef and Ahmad Chatha; Formal analysis, Qudrat Ullah; Investigation, Maria Lateef and Muhammad Hassan; Methodology, Saima Mustafa, Abdul Qadeer and Muhammad Hassan; Software, Qudrat Ullah and Ahmad Chatha; Supervision, Saima Mustafa; Writing – original draft, Qudrat Ullah, Dalia Fouad, Abdul Qadeer and Maria Lateef; Writing – review & editing, Saima Mustafa, Abdul Qadeer, Muhammad Hassan and Ahmad Chatha

## Appendix A

**Table S1.** Average weight, Fork and Total lengths of fish sampled from the study sites at LB and RB of Punjnad. headworks.

| Fish Species |  | Average weight (g) |  |  | Average Fork Length (inch) |  |  | Average Total Lengths (inch) |  |  |
| --- | --- | --- | --- | --- | --- | --- | --- | --- | --- | --- |
|  |  | Winter | Spring | Summer | Winter | Spring | Summer | Winter | Spring | Summer |
| *C. mrigala* | RB | 270.90 | 256.42 | 261.49 | 27.26 | 16.87 | 21.1 | 29.495 | 19.98 | 23.73 |
|  | LB | 281.02 | 280.74 | 262.94 | 32.91 | 32.83 | 28.35 | 46.31 | 35.68 | 31.61 |
| *W. attu* | RB | 263.92 | 277.44 | 270.87 | 42.57 | 57.93 | 51.72 | 45.6 | 60.96 | 54.69 |
|  | LB | 195.82 | 210.33 | 208.22 | 21.64 | 25.53 | 25.35 | 24.69 | 28.54 | 28.02 |
| *R. rita* | RB | 208.10 | 226.18 | 223.37 | 37.52 | 45.31 | 43.64 | 40.48 | 48.32 | 46.67 |
|  | LB | 181.13 | 185.55 | 194.83 | 27.42 | 29.23 | 29.92 | 30.38 | 32.31 | 33.06 |
| *M. armatus* | RB | 274.67 | 283.89 | 291.99 | 33.41 | 38.26 | 42.44 | 41.1 | 41.27 | 45.46 |
|  | LB | 328.68 | 329.05 | 320.99 | 51.47 | 53.91 | 45.71 | 53.44 | 56.05 | 47.71 |
| *S. sarwari* | RB | 291.77 | 295.72 | 287.26 | 40.08 | 43.59 | 34.27 | 47.13 | 50.61 | 41.26 |
|  | LB | 475.65 | 394.60 | 352.65 | 58.56 | 51.55 | 48.32 | 65.53 | 58.55 | 55.69 |

† RB = right bank of Punjnad headworks; LB = left bank of Punjnad headworks

